# Peripheral Exercise Transcriptomes Do Not Converge on Glutamatergic or Neuroinflammatory Signatures: A Cross-Platform Meta-Analysis

**DOI:** 10.64898/2026.10.27.762209

**Authors:** Amrita Balajee, Monazza Shahab

## Abstract

Executive dysfunction following physical overtraining has been linked to glutamatergic dysregulation in the lateral prefrontal cortex (LPFC), but whether this central mechanism is reflected in peripheral gene expression is unknown. This study tested whether peripheral transcriptomic responses to exercise converge on the glutamatergic or neuroinflammatory pathways implicated in LPFC dysfunction. Three publicly available GEO datasets spanning skeletal muscle, whole blood, and circulating neutrophils were analyzed across four exercise groups using limma, with a discovery threshold of nominal p < 0.05 and absolute log2 fold change above 0.5. Differential expression analysis identified 7,311 unique differentially expressed genes (DEGs) across the four contrasts, but overlap between datasets was limited: 7 genes were shared between skeletal muscle and blood, 38 between skeletal muscle and the GSE43856 contrasts, and 90 between blood and the GSE43856 contrasts, with none significant across all three studies. Gene Ontology and KEGG enrichment of the DEG union identified ribonucleoprotein complex biogenesis, ribosome biogenesis, protein folding, and T cell activation as the dominant biological processes; no glutamatergic or neuroinflammatory terms reached significance. Cross-tissue concordance between skeletal muscle and blood was absent (r = 0). Effect sizes among the genes driving the top enriched pathways were dominated by ribosomal proteins, heat shock proteins, and translation initiation factors rather than glutamate receptors or transporters. These findings indicate that peripheral transcriptomic responses to exercise are tissue-specific and dominated by proteostasis, translational, and immune programs rather than glutamatergic signaling. Direct assessment of central nervous system tissue or fluid-based glutamatergic biomarkers may be required to link overtraining-induced fatigue to LPFC dysfunction.

## Introduction

Athletic performance depends not only on physical capacity but on the cognitive systems that govern it (Sun et al., 2025). Athletes must continuously evaluate risk, suppress impulsive responses, and execute rapid strategic decisions under psychological and physiological stress (Ren et al., 2024), making executive function a meaningful, underappreciated determinant of performance.

In soccer players, mental fatigue is associated with reduced intermittent running performance, increased passing and ball-control errors, and decreased shot speed and accuracy (Smith et al., 2016), with subsequent work extending these effects to reduced dribbling accuracy, degraded tactical judgment, and increased ineffective passing under pressure (Trecroci et al., 2020; Gantois et al., 2020). These decrements likely reflect fatigue-induced disruption of goal-directed attention and cognitive control (Sun et al., 2021), and mental-fatigue-related performance deficits appear to be driven by executive-control disruption rather than declines in physical capacity alone (Van Cutsem et al., 2017) together establishing that mental fatigue disrupts the executive-control processes central to adaptive decision-making in competition.

The neurochemical basis of this disruption is converging on glutamatergic dysfunction in the lateral prefrontal cortex (LPFC): prolonged cognitively demanding tasks produce measurable LPFC glutamate accumulation alongside behavioral shifts toward impulsivity and effort avoidance (Wiehler et al., 2022), suggesting that sustained cognitive exertion overwhelms the LPFC’s capacity for glutamate clearance and metabolic regulation, degrading inhibitory control (Möller, 2023). Beyond cortical accumulation, the metabotropic glutamate receptors mGluR1 and mGluR5 maintain behavioral flexibility and adaptive action selection via corticostriatal plasticity and executive-control circuits, and their disruption is associated with impaired cognitive flexibility, perseverative behavior, and deficits in goal-directed decision-making in animal models (Xu et al., 2009). This mechanism extends beyond cognitive fatigue: overtrained mice show reduced mGluR1/5 signaling in the dorsolateral striatum alongside heightened behavioral inflexibility (Paget-Blanc et al., 2025), suggesting that physical overtraining can similarly compromise the glutamatergic circuitry underlying adaptive decision-making, and raising the possibility that exhaustion and overtraining converge on a shared glutamatergic mechanism that erodes executive control.

Despite this emerging picture, the relationship between fatigue-associated glutamatergic alterations and athletic cognitive performance remains incompletely understood. This study addresses that gap through a transcriptomic meta-analysis of publicly available GEO/NCBI datasets, testing whether molecular signatures of intense exercise and fatigue map onto the glutamatergic pathways implicated in LPFC dysfunction and executive impairment links with implications both for understanding how overtraining degrades performance and for informing strategies to prevent it.

To address this, three publicly available GEO datasets were integrated: skeletal muscle profiles across varying exercise intensities (GSE48278), blood-based responses to high-intensity interval training (GSE10540), and circulating neutrophil and skeletal muscle changes following prolonged endurance exercise (GSE43856). These peripheral-tissue datasets serve as accessible proxies for the systemic molecular response to exercise and overtraining, since direct transcriptomic profiling of brain tissue is largely unobtainable in living humans. This meta-analysis integrates these independent datasets to determine whether they converge on a shared molecular signature; consistent implication of glutamatergic or neuroinflammatory pathways across peripheral tissues would suggest accessible biomarkers for centrally-mediated cognitive decline associated with overtraining.

## Methods

### Dataset selection and inclusion criteria

Three publicly available gene expression datasets were retrieved from the NCBI Gene Expression Omnibus (GEO), selected to represent distinct exercise modalities, tissues, and timescales of transcriptomic response. GSE48278 contained skeletal muscle (vastus lateralis) profiles from the STRRIDE program, sampled before and after exercise training (114 samples; Affymetrix Human Genome U133 Plus 2.0, GPL570; Barberio et al., 2016). GSE10540 contained whole-blood profiles from metabolic syndrome patients sampled before and after 16 weeks of aerobic interval training (10 samples; ABI Human Genome Survey Microarray, GPL2986; Bye et al., 2009). GSE43856 contained circulating neutrophil (PMN) and skeletal muscle profiles from healthy endurance-trained males sampled at baseline and at 3, 48, and 96 hours after an experimental exercise trial (64 samples; Illumina HumanHT-12 V4.0, GPL10558; Neubauer et al., 2013, 2014).

The three datasets span distinct microarray platforms; cross-platform integration was therefore performed at the level of statistical results rather than raw expression values, consistent with established approaches for cross-platform transcriptomic meta-analysis.

### Data retrieval and preprocessing

Series matrix files were obtained from GEO and imported into R (v4.6.0) using GEOquery (v2.80.0) and extracted as ExpressionSet objects. Each dataset’s distribution was inspected by per-sample boxplot and density plot; a log2 transformation was applied where maximum intensity exceeded 100, indicating a linear scale. GSE48278 and GSE10540 were log2 transformed; GSE43856 was already log2-scaled and left untransformed. No additional normalization was applied, as each dataset had already been normalized by the original submitters using platform-appropriate methods. Probes with zero variance across samples were removed prior to differential expression analysis.

### Differential expression analysis

Differential expression was assessed using limma (v3.68.3) with empirical Bayes moderation. For each dataset, a linear model was fit and contrasts defined according to experimental design. GSE48278 used a paired design comparing post-versus pre-training skeletal muscle, with subject as a blocking factor (∼subject + group). GSE10540 used an unpaired design because the paired model exhausted the residual degrees of freedom at this sample size (n = 5 pairs); results from this dataset are treated as exploratory throughout. GSE43856 skeletal muscle and neutrophil samples were analyzed separately, each with a paired design comparing the 3-hour post-exercise state to baseline (subject identity extracted from the sample characteristics field).

Probe-level test statistics, log2 fold changes, moderated t-statistics, and p-values were extracted for all probes, with gene symbols obtained from platform feature data. Because the three datasets differ substantially in sample size, platform, and biological question, a nominal p < 0.05 combined with an absolute log2 fold change greater than 0.5 was used as the discovery threshold across all four contrasts, consistent with the analytical approach of the original publications; all results are designated exploratory. Multiple-testing correction was not applied across probes, as the sample sizes of GSE48278 (n = 57 pairs) and GSE10540 (n = 5 pairs) render Benjamini-Hochberg false discovery rate control prohibitively conservative across 54,675 and 32,878 probes, respectively, eliminating all signal.

### Cross-dataset comparison

Overlap of significant DEGs across datasets was assessed by pairwise and three-way intersection of significant gene sets. Cross-tissue concordance was quantified as the Pearson correlation between gene-level log2 fold changes in skeletal muscle (GSE48278) and blood (GSE10540), with fold changes averaged across probes per gene prior to correlation.

### Pathway enrichment analysis

Functional enrichment was performed on the union of all significant DEGs across the four contrasts (7,311 unique genes) using clusterProfiler, with gene symbols mapped to Entrez identifiers via org.Hs.eg.db. Gene Ontology Biological Process (GO BP) terms and Kyoto Encyclopedia of Genes and Genomes (KEGG) pathways were tested for over-representation against a background of all genes assessed across the four contrasts; enrichment p-values were Benjamini-Hochberg adjusted, and terms/pathways with adjusted p < 0.05 were retained.

### Convergence analysis

Genes driving the top five enriched GO BP pathways were extracted from the clusterProfiler results, and the subset also significant in at least one dataset was carried forward for convergence analysis. For each gene, the log2 fold change and 95% confidence interval were derived from the moderated t-statistic (SE = logFC / t) across all four contrasts; effect size, direction of change, and the proportion of datasets reaching nominal significance were summarized in a forest plot and a convergence bubble plot.

### Statistical analysis and software

All analyses were performed in R (v4.6.0). Differential expression used limma (v3.68.3) with empirical Bayes moderation. Pathway enrichment used clusterProfiler with org.Hs.eg.db. Figures were generated using ggplot2, pheatmap, cowplot, and VennDiagram.

## Results

### Exercise induces transcriptomic responses across peripheral tissues

#### Included datasets summary

The included datasets represented distinct tissues, exercise paradigms, and transcriptomic platforms (Figure 1A). GSE48278 examined skeletal muscle before and after exercise training and identified 239 DEGs; GSE10540 assessed blood before and after HIIT and identified 1,145 DEGs; GSE43856 measured acute responses three hours after exercise in both neutrophils and skeletal muscle, identifying 3,941 and 2,711 DEGs, respectively. The number of exercise-responsive genes varied substantially across datasets, consistent with differences in tissue type, study design, and experimental context. Volcano plots (Figure 1B) showed substantial variation in the magnitude and direction of transcriptional change across datasets: relatively few genes exceeded threshold in GSE48278 skeletal muscle, whereas the acute GSE43856 contrasts showed more extensive remodeling, with strong upregulation of *PPARGC1A, NR4A3, ABRA*, and *MAFF* in skeletal muscle and extensive immune- and signaling-related changes in neutrophils. These findings indicate that exercise induces robust but context-dependent transcriptional responses that vary substantially across tissue, exercise paradigm, and sampling time point.

**Table 1.** Characteristics of the datasets and exercise contrasts included in the meta-analysis.

| Dataset | Reference | Tissue | Platform | n | Contrast | DEGs |
|---|---|---|---|---|---|---|
| GSE48278 | Barberio et al., 2016 | Skeletal muscle | Affymetrix HG-U133 Plus 2.0 | 114 | Post vs. pre training | 239 |
| GSE10540 | Bye et al., 2009 | Whole blood | ABI Human Genome Survey | 10 | Post vs. pre HIIT | 1,145 |
| GSE43856 | Neubauer et al., 2013 | Neutrophils | Illumina HumanHT-12 V4.0 | 16 | 3 h post vs. baseline | 3,941 |
| GSE43856 | Neubauer et al., 2014 | Skeletal muscle | Illumina HumanHT-12 V4.0 | 16 | 3 h post vs. baseline | 2,711 |

Note: n, samples analyzed per contrast; DEGs, differentially expressed genes at nominal p < 0.05 and absolute log2 fold change > 0.5; HIIT, high-intensity interval training. Neutrophil and skeletal muscle samples from GSE43856 were analyzed as separate contrasts.

**Figure 1.**
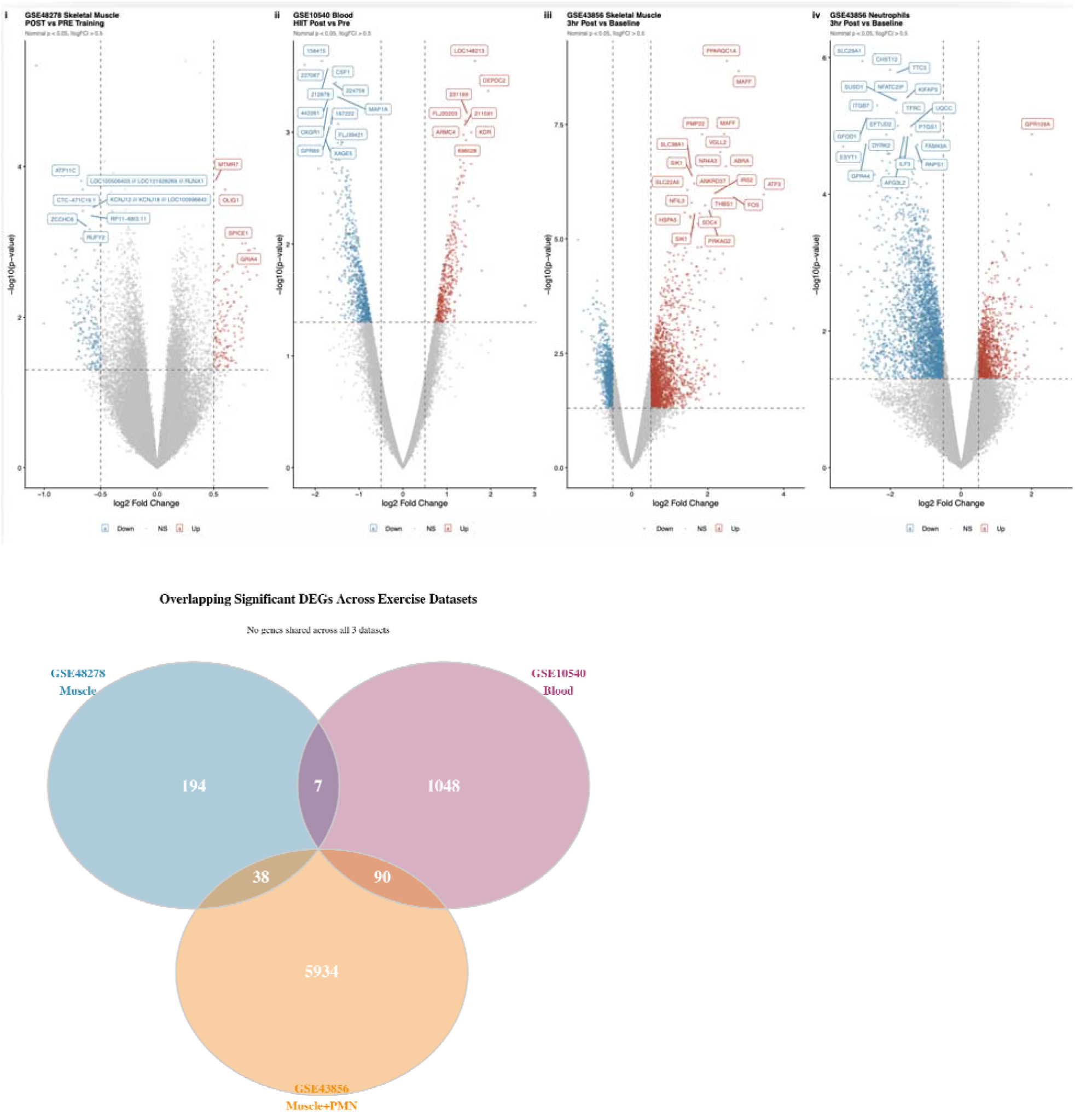
Study design and differential expression across three exercise transcriptome datasets. (A) Summary of the three GEO datasets included in the meta-analysis, listing accession identifier, first author, publication year, tissue, microarray platform, number of samples analyzed, experimental contrast, and number of significant DEGs at nominal p < 0.05 and absolute log2 fold change above 0.5. GSE48278 (skeletal muscle, STRRIDE program) yielded 239 DEGs; GSE10540 (whole blood, aerobic interval training in metabolic syndrome) yielded 1,145 DEGs; GSE43856 neutrophils yielded 3,941 DEGs and GSE43856 skeletal muscle yielded 2,711 DEGs at 3 hours post-exercise versus baseline. (B) Volcano plots of differential expression across the four contrasts: (i) GSE48278 skeletal muscle, POST versus PRE training; (ii) GSE10540 blood, HIIT post versus pre; (iii) GSE43856 skeletal muscle, 3 hours post-exercise versus baseline; (iv) GSE43856 neutrophils, 3 hours post-exercise versus baseline. The x-axis shows log2 fold change and the y-axis shows negative log10 nominal p-value; red points indicate upregulated genes, blue points indicate downregulated genes, and grey points indicate non-significant genes; dashed lines mark the nominal p-value threshold of 0.05 and the fold-change threshold of 0.5; the 20 most significant genes per panel are labeled. GSE10540 and the GSE43856 neutrophil contrast are underpowered; results are exploratory. (C) Venn diagram of overlapping significant DEGs across datasets: three-way overlap of GSE48278 skeletal muscle, GSE10540 blood, and GSE43856 combined muscle and neutrophil. Seven genes were shared between GSE48278 and GSE10540, 38 between GSE48278 and GSE43856, and 90 between GSE10540 and GSE43856. No genes were significant across all three datasets.

#### Limited overlap of differentially expressed genes across datasets

To determine whether exercise-responsive transcriptional programs converged across tissues and experimental conditions, overlap among significant DEGs was assessed. Despite identifying 7,311 unique DEGs across all four contrasts, overlap between datasets was limited (Figure 1C): 7 genes were shared between GSE48278 skeletal muscle and GSE10540 blood, 38 between GSE48278 and GSE43856, and 90 between GSE10540 and GSE43856, with no genes significant across all three datasets. Overlapping genes included a mixture of immune-regulatory, signaling, and structural genes, including *ARG1, ANXA1, THBS1*, and *GRIA4*. The absence of universally shared DEGs indicates that peripheral transcriptomic responses to exercise are highly tissue-specific and strongly influenced by exercise type and physiological context.

#### Functional enrichment identifies proteostasis, translation, and immune activation pathways

Gene Ontology (GO) and KEGG pathway analyses were performed on the union of all significant DEGs. GO Biological Process analysis revealed significant enrichment of ribonucleoprotein complex biogenesis, ribosome biogenesis, cytoplasmic translation, protein folding, rRNA processing, and RNA transport, along with immune processes including T-cell activation, leukocyte activation, and positive regulation of lymphocyte activation (Figure 2A); no glutamatergic signaling pathways or neuroinflammatory GO terms reached significance. KEGG pathway analysis similarly demonstrated enrichment of pathways associated with ribosomal function, oxidative phosphorylation, protein processing in the endoplasmic reticulum, spliceosome activity, carbon metabolism, and cytoskeletal organization (Figure 2B). Neurodegeneration-associated pathways, including Parkinson’s disease, were also enriched, but primarily reflected genes involved in mitochondrial function, oxidative stress responses, and proteostasis rather than disease-specific mechanisms. These findings indicate that exercise-responsive peripheral transcriptomes are dominated by translational, metabolic, and immune-regulatory processes rather than glutamatergic signaling.

**Figure 2.**
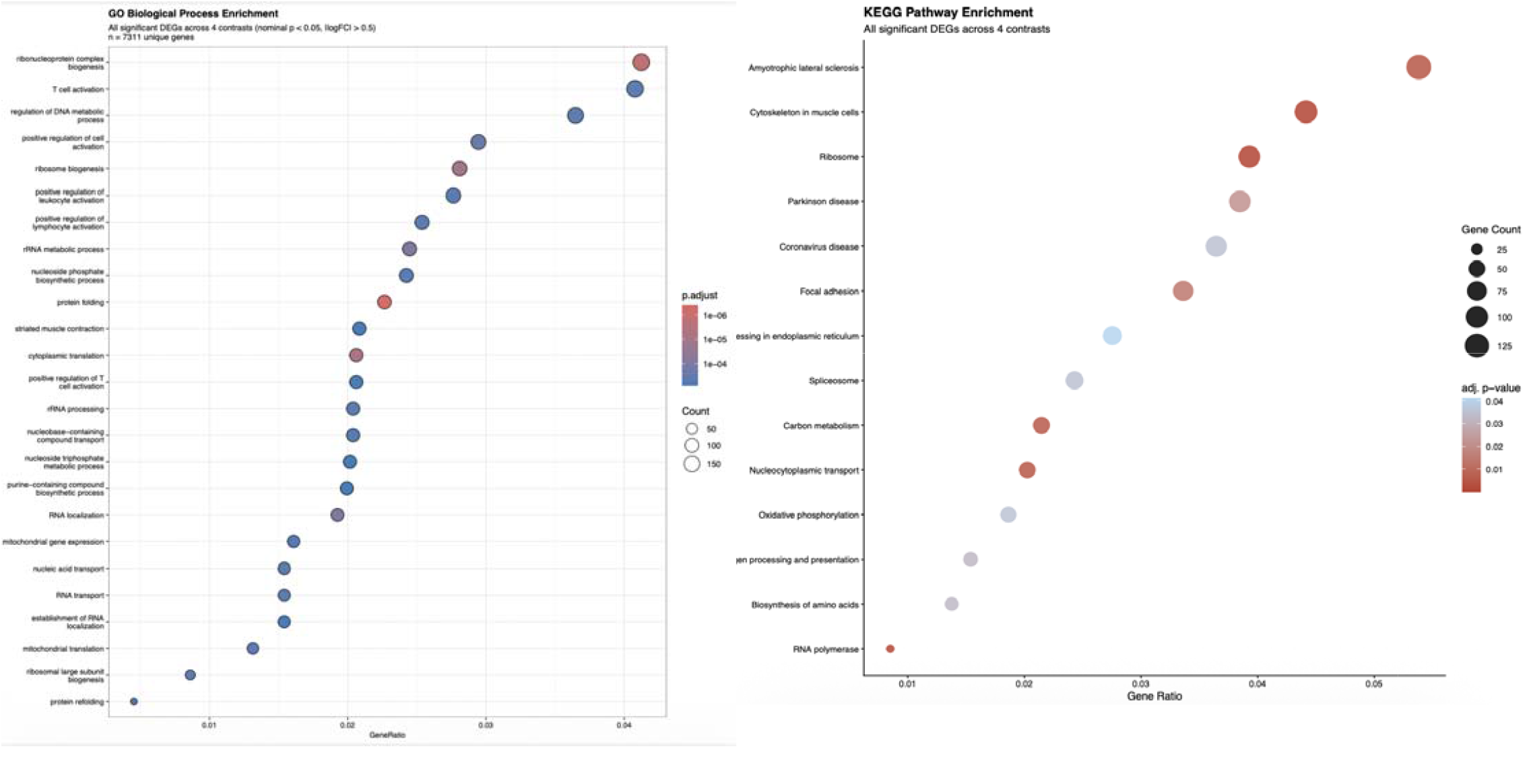
Functional enrichment of exercise-responsive genes. (A) Gene Ontology Biological Process enrichment: dot plot of the 25 most significantly enriched GO Biological Process terms across the union of 7,311 unique significant DEGs from all four contrasts. Point size indicates the number of genes mapping to each term; color indicates adjusted p-value (Benjamini-Hochberg). The most significantly enriched terms include ribonucleoprotein complex biogenesis, T cell activation, ribosome biogenesis, protein folding, striated muscle contraction, and cytoplasmic translation; no glutamatergic or neuroinflammatory terms reached significance; the dominant biological signal reflects proteostasis, translational machinery, and immune cell activation. (B) KEGG pathway enrichment: bubble plot of KEGG pathways enriched in the union of significant DEGs across all four contrasts. The x-axis shows the gene ratio (enriched genes divided by total pathway genes); point size indicates the number of enriched genes per pathway; color indicates adjusted p-value. The most significantly enriched pathways include Ribosome, Cytoskeleton in muscle cells, Amyotrophic lateral sclerosis, Parkinson disease, Oxidative phosphorylation, Protein processing in endoplasmic reticulum, Spliceosome, and Carbon metabolism; enrichment of the neurodegeneration-associated pathways reflects shared gene sets involving mitochondrial function, oxidative stress, and proteostasis rather than disease-specific mechanisms.

#### Genes shared across studies show weak cross-tissue concordance

To further evaluate transcriptomic convergence, the top 50 genes significant in at least two datasets were examine (Figure 3). Although many genes, including *ARG1, ANXA1*, and *THBS1*, along with signaling and cytoskeletal genes, appeared repeatedly across datasets, their expression patterns were not consistent across tissues: genes upregulated in one tissue were frequently unchanged or downregulated in another, indicating limited cross-tissue reproducibility even among overlapping DEGs. Comparison of gene-level fold changes between skeletal muscle (GSE48278) and blood (GSE10540) revealed almost no global correlation (Figure 4): the Pearson correlatio coefficient was approximately zero, indicating that transcriptional responses observed in skeletal muscle were largely independent of those observed in blood.

**Figure 3.**
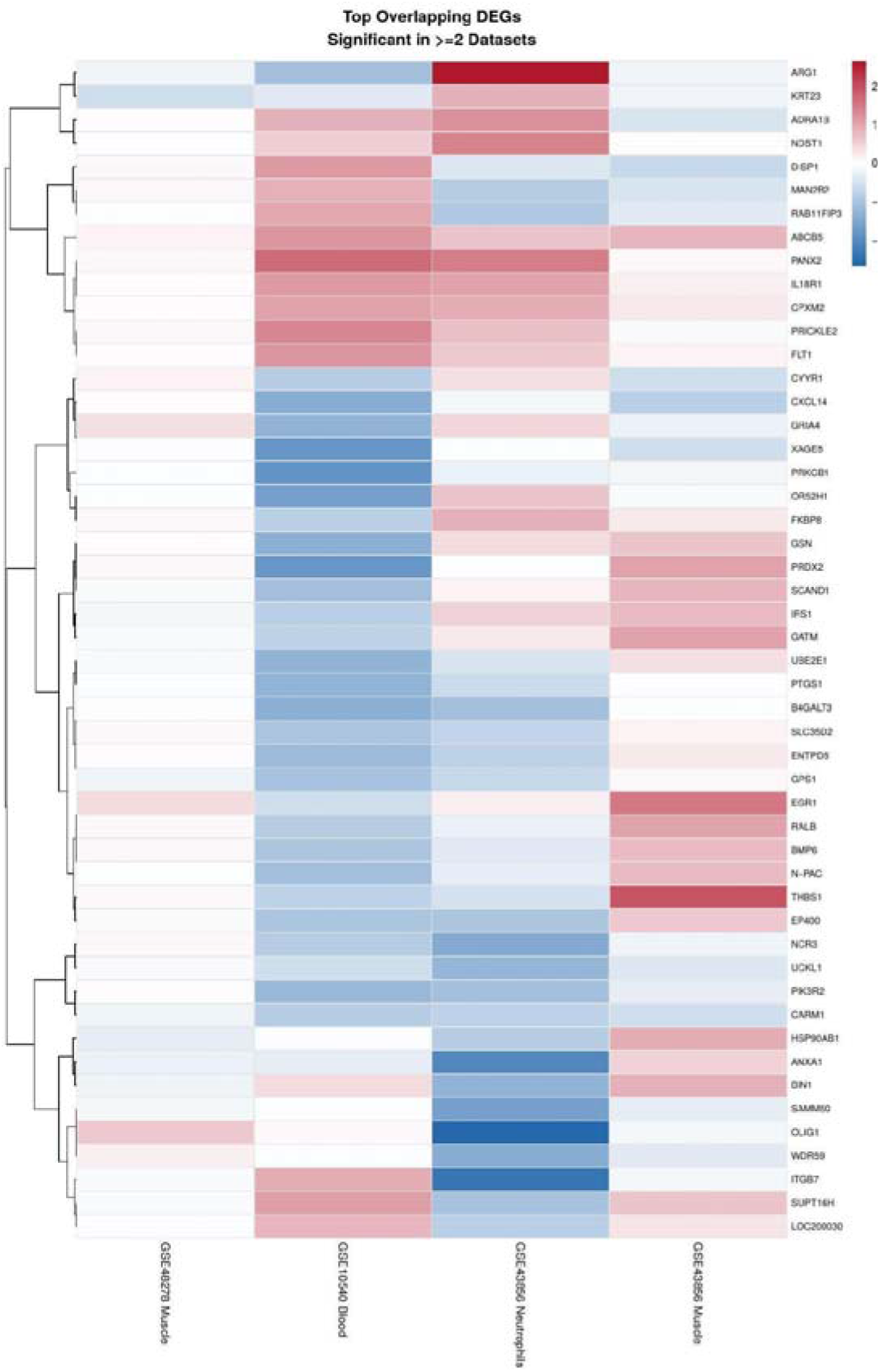
Heatmap of top overlapping DEGs across exercise datasets. Heatmap of log2 fold change for the top 50 DEGs significant in at least two datasets, ranked by mean absolute fold change. Rows represent genes and columns represent the four contrasts (GSE48278 skeletal muscle, GSE10540 blood, GSE43856 neutrophils, GSE43856 skeletal muscle). Red indicates increased expression and blue indicates decreased expression relative to the comparison condition; missing values (grey) indicate the gene was not significant or not detected in that contrast. Overlapping DEGs include immune-regulatory genes such as *ARG1, ANXA1*, and *THBS1*, alongside cytoskeletal and signaling genes. The pattern of direction and magnitude is inconsistent across tissues, indicating limited cross-tissue concordance even among shared DEGs.

**Figure 4.**
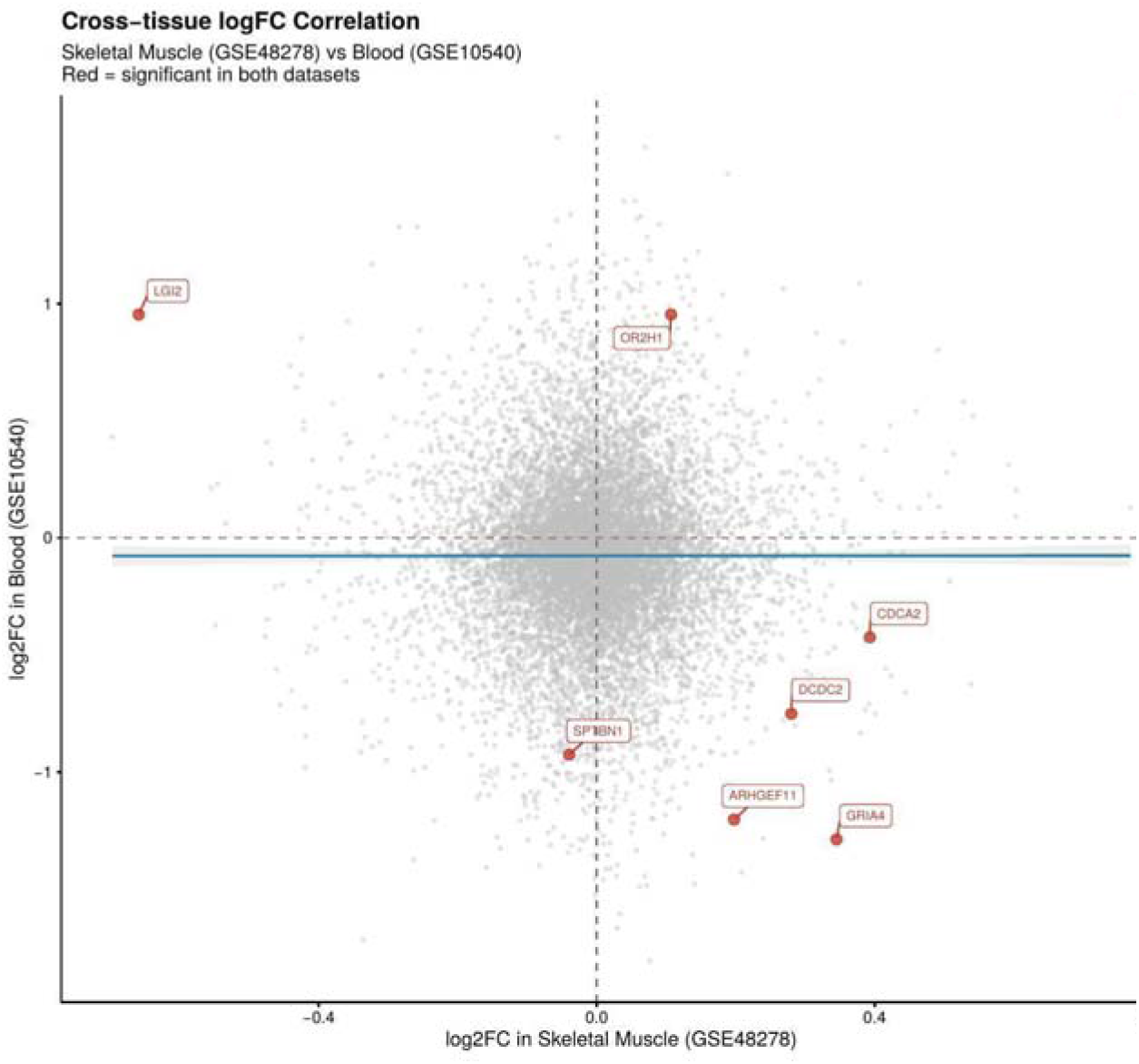
Cross-tissue log2 fold-change correlation between skeletal muscle and blood. Scatter plot of gene-level log2 fold change in skeletal muscle (GSE48278, x-axis) against log2 fold change in blood (GSE10540, y-axis) for all genes detected in both datasets. Each point represents one gene; red points indicate genes significant in both datasets simultaneously (n = 7). The Pearson correlation coefficient between muscle and blood fold changes was r = 0, indicating no linear concordance between the transcriptomic responses of skeletal muscle and peripheral blood to exercise. Labeled genes include *GRIA4, ARG1, FLT1*, and *GSN*. This result indicates that blood transcriptomics cannot serve as a proxy for skeletal muscle transcriptomics in the context of exercise training.

#### Cross-dataset convergence of top GO pathway genes

To assess whether genes driving the top enriched GO pathways showed consistent responses across datasets, effect sizes were compared across all four contrasts using a forest plot (Figure 5A). These genes were largely involved in ribosomal function, translation, protein folding, and RNA processing, including *RPL*/*RPS* family members, *HSPA8, HSP90B1, EIF3E*, and *SNRNP70*. Significant effects were most prominent in the GSE43856 neutrophil and skeletal muscle contrasts, where many genes were downregulated, whereas GSE48278 skeletal muscle and GSE10540 bloo showed smaller and less consistent changes. Mean fold change, effect magnitude, direction, and cross-dataset significance rate for these genes are further summarized in a convergence bubble plot (Figure 5B), which shows that the predominant signal is downregulation, with the highest convergence for ribosomal and heat shock protein genes across the acute GSE43856 contrasts. Overall, the variation in effect direction and magnitude across datasets suggests limited convergence even among genes contributing to the most strongly enriched pathways.

**Figure 5.**
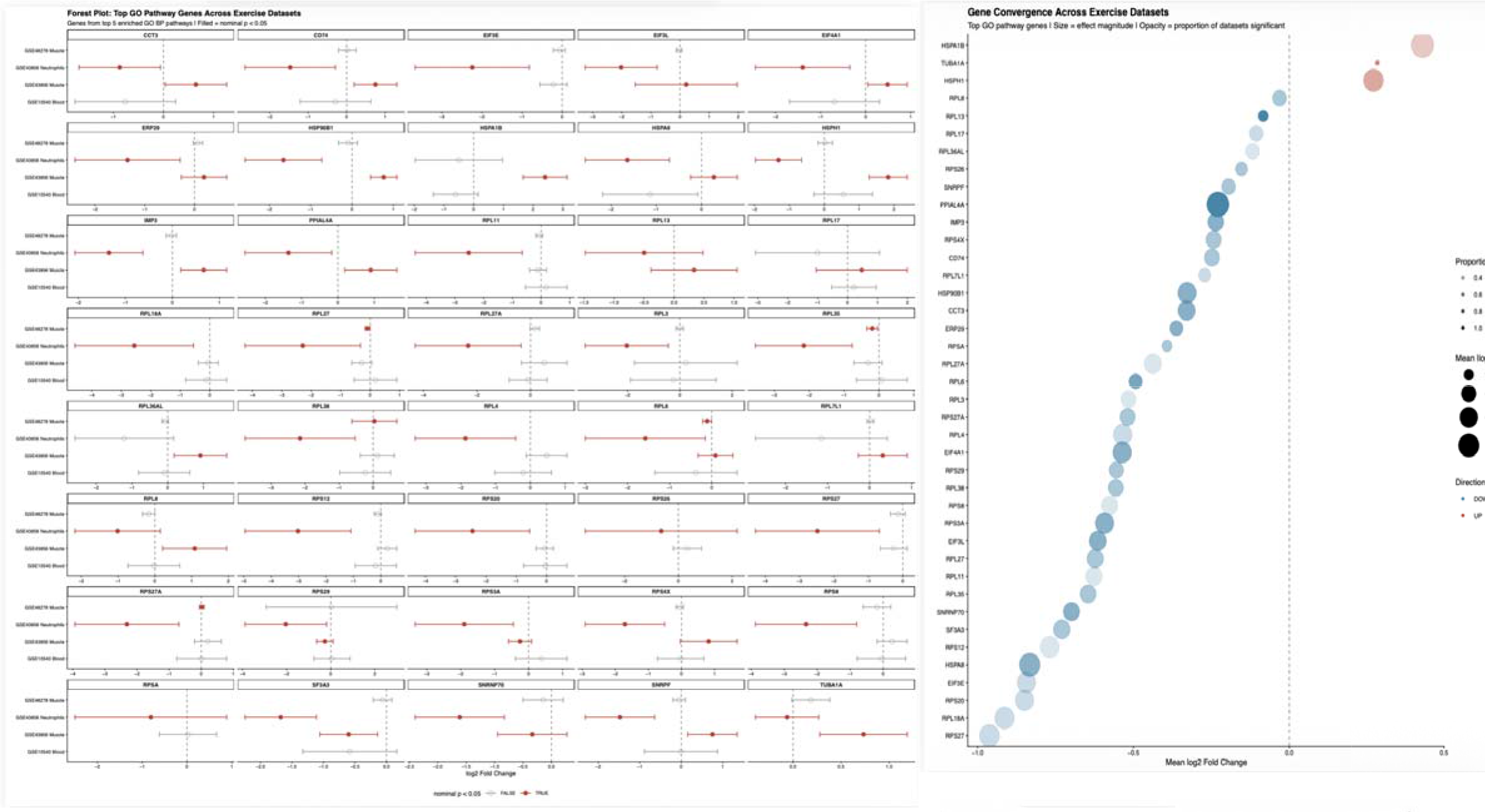
Cross-dataset convergence of top GO pathway genes. (A) Forest plot showing the log2 fold change and 95% confidence interval for the top 40 genes driving the five most significantly enriched GO Biological Process pathways, displayed across the four experimental contrasts. Points represent the mean log2 fold change for that gene in each dataset; horizontal bars represent the 95% confidence interval derived from the moderated t-statistic; filled circles indicate nominal p < 0.05, open circles indicate non-significant results. Genes are dominated by ribosomal proteins (*RPL* and *RPS* family members), heat shock proteins (*HSPA8, HSPH1, HSP90B1*), translation initiation factors (*EIF3E, EIF3L, EIF4A1*), and RNA processing components (*SNRNP70, SF3A3*); most genes show significant downregulation in the GSE43856 skeletal muscle and neutrophil contrasts, with smaller and less consistent effects in GSE48278 and GSE10540. (B) Convergence bubble plot summarizing the mean log2 fold change, effect magnitude, direction of change, and cross-dataset significance rate for each of the top GO pathway genes. The x-axis shows the mean log2 fold change across all four contrasts; point size reflects the mean absolute fold change; color indicates direction (red = upregulated, blue = downregulated on average); opacity reflects the proportion of the four datasets in which the gene reached nominal significance (p < 0.05). The predominant signal is downregulation, with the highest convergence observed for ribosomal and heat shock protein genes across the acute exercise contrasts of GSE43856.

## Discussion

This study examined whether exercise-responsive transcriptional changes converge across peripheral tissues, with particular focus on glutamatergic or neuroinflammatory signaling. Although thousands of differentially expressed genes were identified across skeletal muscle, blood, and neutrophils, overlap between datasets was limited, and no gene was significantly altered across all three datasets. Instead, the strongest enriched biological processes were related to ribonucleoprotein and ribosome biogenesis, translation, protein folding, muscle contraction, and immune activation; the glutamatergic and neuroinflammatory gene ontology pathways showed no significance. These results suggest that peripheral transcriptional responses to exercise are highly dependent on tissue type and experimental context rather than reflecting a single conserved molecular program.

The dominant enrichment of pathways related to translation, ribosomal activity, and protein folding suggests that exercise strongly alters cellular processes involved in protein synthesis and proteostasis. However, even these genes did not show uniform responses across datasets: the most significant changes were in the acute GSE43856 skeletal muscle and neutrophil contrasts, while effects in GSE48278 skeletal muscle and GSE10540 blood were generally smaller and less consistent. This variability shows that even the most strongly represented exercise-responsive pathways may differ substantially according to tissue and timing of measurement.

The limited overlap among DEGs also supports the importance of tissue specificity. Only seven genes were shared between GSE48278 skeletal muscle and GSE10540 blood, 38 between GSE48278 and GSE43856, and 90 between GSE10540 and GSE43856, with no genes shared across all three datasets, and fold changes in skeletal muscle and blood showed minimal linear concordance (Pearson r ≈ 0), indicating that shared variance between the two tissues’ transcriptomic responses was negligible. This pattern is consistent with the expectation that skeletal muscle, blood, and neutrophils differ fundamentally in baseline transcriptional programs, turnover rates, and exposure to exercise-derived signals, such that a shared molecular response was never assured a priori. Rather than indicating a failure of convergence, this tissue-specific patterning points to protein synthesis and proteostatic regulation as a common cellular currency of the exercise response, even where the specific genes and direction of change differ across tissues. These findings further suggest that blood transcriptional responses should not be assumed to directly reflect those occurring in skeletal muscle, and more broadly highlight the difficulty of using a single peripheral tissue as a proxy for systemic or centrally mediated responses to exercise.

Several features of the included studies may explain the limited convergence observed. The datasets differed in tissue, participant population, exercise paradigm, sampling time, and microarray platform: GSE48278 examined skeletal muscle following an exercise-training intervention, GSE10540 assessed blood after aerobic interval training in individuals with metabolic syndrome, and GSE43856 measured skeletal muscle and neutrophils three hours after an acute exercise trial in endurance-trained males, probably capturing different stages of the physiological response to exercise, ranging from acute cellular stress to longer-term adaptation. The absence of a shared glutamatergic or neuroinflammatory signature in these datasets does not necessarily exclude the involvement of these pathways in exercise-related fatigue, but suggests that such mechanisms were not detectably conserved in the peripheral transcriptomic datasets examined here. Differential expression was assessed using a p < 0.05 and an absolute log2 fold-change threshold greater than 0.5, and multiple-testing correction was not applied at the individual-probe level; future work could address this using larger, prospectively designed cohorts with standardized exercise protocols, matched sampling times, and simultaneous measurement of multiple tissues, alongside studies directly relating peripheral transcriptomic changes to central measures of fatigue or glutamatergic function, which may be particularly important for determining whether peripheral biomarkers can capture molecular processes associated with exercise-induced cognitive fatigue.

The enrichment of protein synthesis and proteostasis-related pathways is also notable given the established role of proteostatic decline as a hallmark of cellular aging. While the present acute post-exercise contrasts capture transient downregulation of ribosomal and chaperone-related genes rather than durable proteostatic enhancement, longitudinal designs capturing repeated exercise bouts over weeks to months may help determine whether cumulative exercise exposure shifts these same pathways toward improved proteostatic capacity, a question with direct relevance to exercise-based interventions for cellular aging and healthspan. This dominance of proteostasis and mitochondrial-adjacent signaling also aligns with a broader literature implicating cellular energetic and protein-quality-control stress in central fatigue states: mitochondrial dysfunction has been repeatedly identified as a contributing mechanism in myalgic encephalomyelitis/chronic fatigue syndrome (Morris and Maes, 2014; Holden et al., 2020), and a growing body of work extends this mitochondrial allostatic load framework to chronic emotional stress and burnout more broadly (Venkatesan et al., 2026). These conditions share phenomenological overlap with athletic overtraining, including persistent fatigue unrelieved by rest and cognitive impairment, raising the possibility that the peripheral molecular signature identified here reflects a more general cellular stress response associated with fatigue states, rather than a pathway specific to glutamatergic or neuroinflammatory mechanisms. Testing whether this proteostasis-dominant signature is shared across overtraining, chronic fatigue syndrome, and burnout populations may clarify whether a common peripheral biomarker profile underlies fatigue-related cognitive impairment across these conditions.

The enrichment of neurodegeneration-associated KEGG pathways, including amyotrophic lateral sclerosis and Parkinson’s disease, most likely reflects the substantial overlap in mitochondrial function, oxidative phosphorylation, and ribosomal gene sets that these pathway annotations share with the exercise-responsive gene set, rather than a direct mechanistic link between exercise and neurodegenerative disease risk. KEGG neurodegeneration pathway terms frequently appear as top hits in transcriptomic studies of clinically unrelated conditions, driven by their heavy reliance on housekeeping oxidative phosphorylation and ribosomal genes rather than disease-specific machinery (Zhao et al., 2025). At the same time, whole-blood transcriptomic studies in ALS itself report genuine enrichment of these same pathways with disease relevance (Zhao et al., 2025), so this overlap is not necessarily uninformative. This raises a testable question for future work: whether the direction and magnitude of change in these shared genes differs meaningfully between healthy exercise-induced stress and pathological neurodegenerative states, which would help clarify whether this convergence is coincidental or mechanistically informative.

The enrichment of T-cell activation and related immune pathways, particularly in the neutrophil contrast, may also be relevant to the inflammaging literature, which links chronic low-grade immune activation to the aging process and to age-related decline in tissue and immune function (Franceschi et al., 2000; Ferrucci and Fabbri, 2018). While the immune signal identified here likely reflects an acute post-exercise response rather than chronic inflammation, acute and chronic immune activation share overlapping regulatory pathways, and T-cell-mediated inflammatory processes have been specifically implicated in age-related inflammatory decline (Mekker et al., 2012). Whether repeated exercise-induced immune activation of the kind identified here contributes cumulatively to an inflammaging-like phenotype in overtrained individuals is a question that longitudinal immune profiling could address directly.

## Conclusion

This study integrated three independent exercise transcriptome datasets spanning skeletal muscle, whole blood, and circulating neutrophils. Peripheral transcriptomic responses to exercise were tissue-specific and dominated by proteostasis, translational, and immune programs rather than glutamatergic signaling. Direct assessment of central nervous system tissue or fluid-based glutamatergic biomarkers may be required to link overtraining-induced fatigue to LPFC dysfunction.

## Data Availability

All datasets analyzed in this study are publicly available from the NCBI Gene Expression Omnibus under accession numbers GSE48278, GSE10540, and GSE43856 (https://www.ncbi.nlm.nih.gov/geo/). All analysis code is openly available at https://github.com/Monazzashahab/lpfc-transcriptomic-meta-analysis.

## Author Contributions

A. Balajee: Conceptualization, Investigation, Data curation, Formal analysis, Visualization, Writing (original draft).

M. Shahab: Conceptualization, Methodology, Supervision, Formal analysis, Validation, Visualization, Writing (review and editing). Both authors read and approved the final manuscript.

## Acknowledgments

The authors thank STEM Bridge Advisors for providing software and subscription resources that supported this work. The authors also thank the investigators who made the GSE48278, GSE10540, and GSE43856 datasets publicly available through the NCBI Gene Expression Omnibus.

## STATEMENT OF AI USAGE

Claude AI used to follow the formatting guidelines for JBCB journals.

